# A novel 5-HT_2A_ receptor-linked signalling pathway involving NAADP-mediated lysosomal Ca^2+^ release

**DOI:** 10.64898/2026.10.29.762736

**Authors:** Aurelija Ippolito, Gary Gilmour, Antony Galione, Grant Churchill, Trevor Sharp

## Abstract

Currently there is major research interest in the therapeutic potential of serotonergic psychedelic drugs across a range of neuropsychiatric disorders^1–3^. Such drugs exert psychedelic and neuroplastic effects, both likely linked to therapeutic outcomes, via agonist action at the 5-HT_2A_ receptor^4–7^. This receptor is assumed to generate downstream intracellular signalling via an interaction with either G_q_ protein coupling and formation of the second messenger inositol trisphosphate (IP_3_), or through triggering the effector β-arrestin2. Central to the IP_3_ pathway is mobilisation of Ca^2+^ stored in the endoplasmic reticulum (ER). Here, we report the unexpected discovery that the 5-HT_2A_ receptor also signals via an additional second messenger, nicotinic acid adenine dinucleotide phosphate (NAADP), which mobilises Ca^2+^ release from lysosomal stores. This effect was revealed through measurement of changes in cellular NAADP levels, pharmacological inhibition of NAADP and direct biosensor measurement of lysosomal Ca^2+^ release and was produced not only by 5-HT but also by a range of psychedelic drugs in a receptor-specific manner. Moreover, genetic ablation of the NAADP pathway reduced markers of both hallucinogenic and neuroplastic effects of psychedelic drug administration in mouse models. These findings reveal the involvement of a novel, therapeutically relevant signalling pathway in psychedelic drug action.

## Introduction

Psychedelic drugs acting on brain neurotransmitter serotonin (5-hydroxytryptamine; 5-HT) receptors are currently under investigation for therapeutic applications in a broad range of psychiatric and neurodegenerative disorders^1–3,8^. The explanation for their mechanism of action across this diversity of diseases is uncertain but their psychedelic effects, as well as their ability to increase different modes of neuroplasticity, have been strongly implicated^9–12^. Of the fourteen 5-HT receptor sub-types in the brain, agonist action specifically at 5-HT_2A_ receptors is thought to trigger both psychedelic and neuroplastic effects of psychedelic drugs^4–7,13–27^. However, a presently unresolved question is which 5-HT_2A_ receptor-mediated downstream signalling pathways underpin the effects of psychedelic drugs and ultimately their therapeutic actions.

5-HT_2A_ receptors are known to couple to a canonical G_q_ protein/phospholipase C-β pathway that increases the formation of the second messengers inositol trisphosphate (IP_3_) and diacylglycerol (DAG) as well as a non-canonical pathway which activates the effector molecule β-arrestin2^28–31^. Several studies have reported evidence of the involvement of G_q_ signalling in psychedelic drug action^17,22,29,32^ although there are conflicting data regarding a possible role of β-arrestin2^33,34^.

Once generated by 5-HT_2A_ receptor activation, IP_3_ binds to IP_3_ receptors, ligand-gated Ca^2+^ release channels on the endoplasmic reticulum (ER), to mobilise Ca^2+^ which then mediates a wide range of neuronal signalling events ^35^. Here we report that, unexpectedly, depletion of ER Ca^2+^ stores (using thapsigargin, a selective inhibitor of ER Ca^2+^ ATPases^36^) did not fully abolish 5-HT_2A_ receptor-mediated Ca^2+^ release, suggesting mobilisation of an additional Ca^2+^ store. On further investigation we discovered that activation of 5-HT_2A_ receptors has the further capacity to generate the formation of another potent Ca^2+^ mobilising second messenger, nicotinic acid adenine dinucleotide phosphate (NAADP)^37^ which specifically releases Ca^2+^ stored in lysosomes via two-pore channels (specifically TPC2). Furthermore, studies in genetically engineered TPC2 null mice suggest that the NAADP/lysosomal pathway plays a role in both psychedelic and neuroplastic effects of psychedelic drugs.

## Results

### 5-HT evokes the release of Ca^2+^ from lysosomal stores

Our previous experiments using neuronal cell lines (rat C6 glioma, human SH-SY5Y neuroblastoma) found that 5-HT evoked a concentration-dependent increase in cytosolic Ca^2+^ mediated solely via the 5-HT_2A_ receptor^28^. Given that the 5-HT_2A_ receptor is G_q_ coupled to phospholipase C and the formation of IP_3_, we predicted that depletion of ER Ca^2+^ stores using thapsigargin (selective inhibitor of ER Ca^2+^ ATPases^36^) would abolish the 5-HT response (Fig. 1A). Unexpectedly, however, in C6 cells pre-treated with thapsigargin (1 μM), 5-HT (0.1 nM - 10 μM) continued to evoke a significant release of Ca^2+^ (Fig. 1B, C; Suppl. Table 1), suggesting that 5-HT mobilised Ca^2+^ from a non-ER source. We replicated the thapsigargin experiment using SH-SY5Y cells transfected with the human 5-HT_2A_ receptor and similarly observed a residual Ca^2+^ response to 5-HT (Fig. 1B; Suppl. Table 2). In contrast, in C6 cells the Ca^2+^ response to ATP (10 nM - 1 mM; likely mediated by G_q_-coupled purinergic P_2Y_ receptors^38^) was completely abolished by thapsigargin (1 μM), indicating specificity of the 5-HT effect (Fig. 1B, D; Suppl. Table 3).

**Figure 1.**
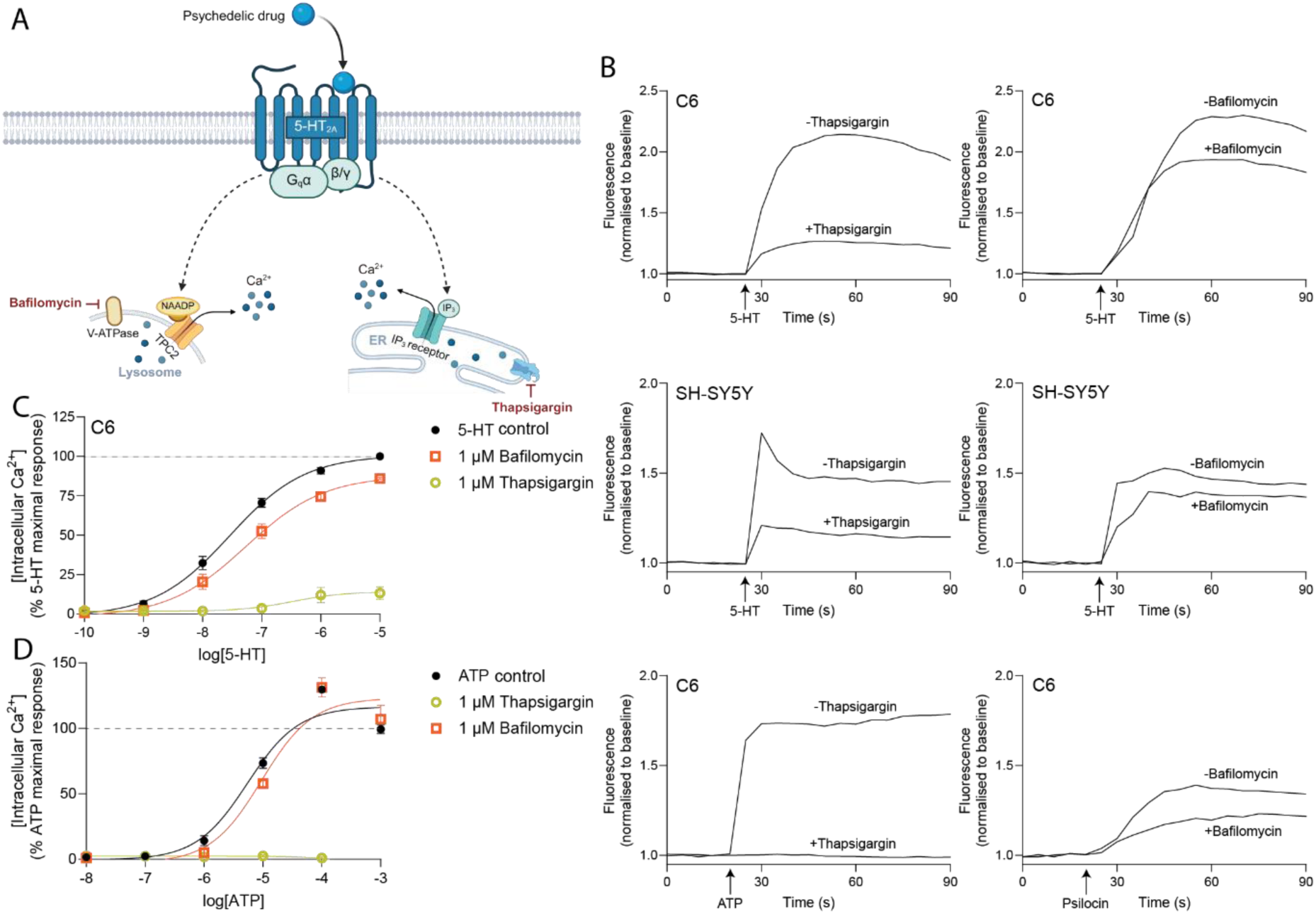
Lysosomal Ca^2+^ contributes to Ca^2+^ responses evoked by the 5-HT_2A_ receptor - use of Ca^2+^ store inhibitors. (A) Schematic representation of links between the 5-HT_2A_ receptor and lysosomal and ER Ca^2+^ stores associated and their inhibitors. (B) Representative raw traces of individual Ca^2+^ responses to 5-HT, ATP or psilocin (10 µM) in C6 or SH-SY5Y cells in the presence of thapsigargin or bafilomycin (1 µM), as specified. (C) Grouped data showing 5-HT concentration-dependent Ca^2+^ responses in C6 cells treated with bafilomycin (p=0.0006) or thapsigargin (p<0.0001, both Extra-sum-of-squares F-test with Dunnett’s test) (1 µM). (D) As with (C) but for ATP. Each point is the mean ± SEM value of five independent experiments performed in duplicates. Responses are relative to 10 µM 5-HT or ATP.

Given these findings, we explored the possibility that 5-HT released Ca^2+^ from a separate Ca^2+^ store, the lysosome (Fig. 1A). To this end, lysosomal Ca^2+^ stores were depleted with bafilomycin (1 μM; selective inhibitor of vacuolar H^+^-ATPases, which abrogates lysosomal Ca^2+^ storage^39,40^) prior to addition of 5-HT or ATP. Bafilomycin reduced the magnitude of the Ca^2+^ response to 5-HT in both C6 and transfected SH-SY5Y cells but had no effect on the response to ATP (Fig. 1B-D; Suppl. Tables 1-3), suggesting a selective mobilisation of lysosomal Ca^2+^ stores by 5-HT response. We have previously shown that the psychedelic drug psilocin evoked a 5-HT_2A_ receptor-mediated Ca^2+^ release^28^, and this response was also reduced by bafilomycin (Fig. 1B; Suppl. Table 4).

### NAADP signalling plays a role in Ca^2+^ response to 5-HT

Given that NAADP releases lysosomal Ca^2+^ (Fig. 2A), we explored the possible involvement of this second messenger in the 5-HT Ca^2+^ response. First, the effect of 5-HT on NAADP synthesis was determined. Compared to vehicle controls, 5-HT (10 μM) caused a significant increase in NAADP levels in C6 cells, and this effect was replicated by the potent and selective 5-HT_2A_ receptor agonist, 25B-NBOMe (10 μM) (Fig. 2B).

**Figure 2.**
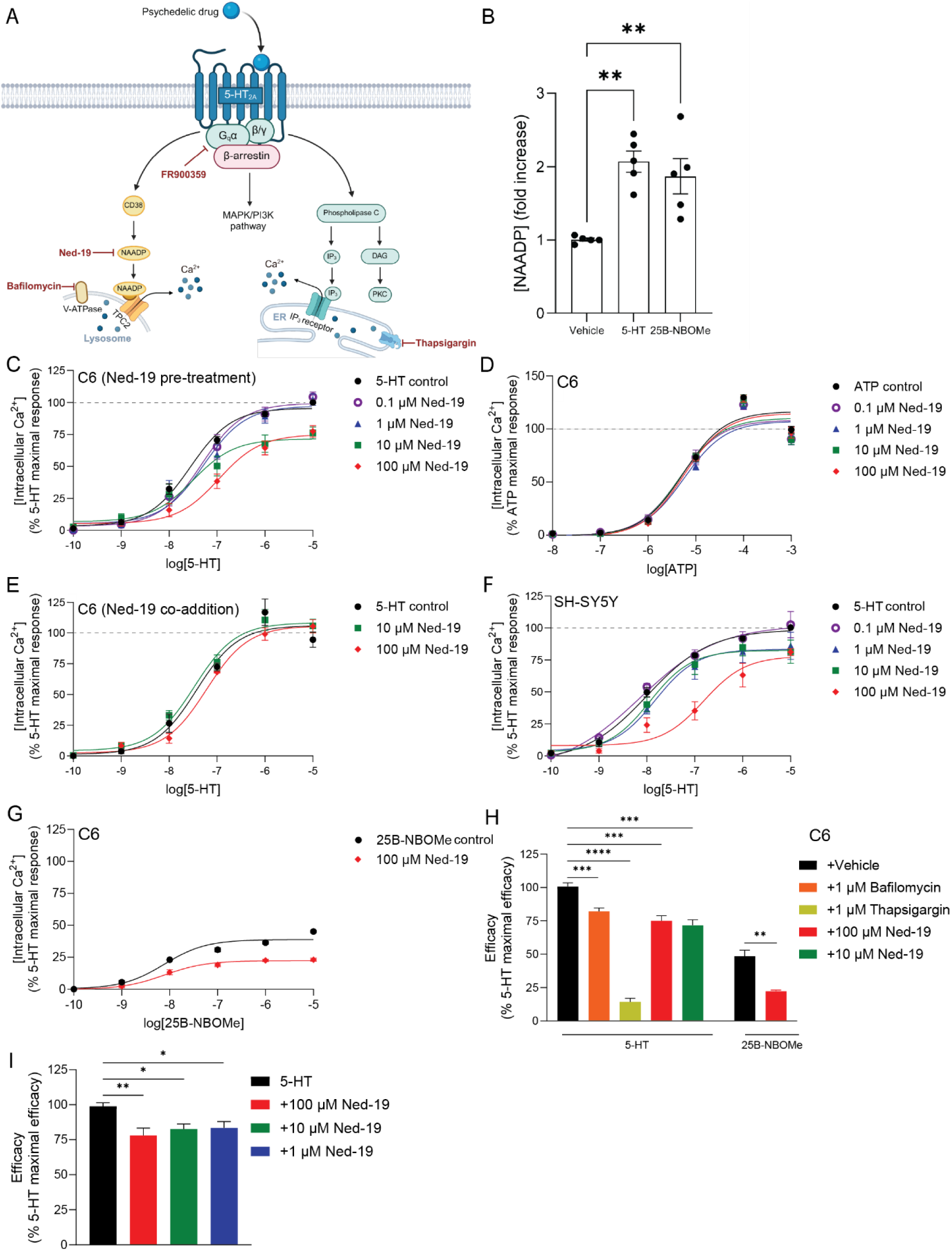
The NAADP/lysosomal pathway contributes to Ca^2+^ responses evoked by the 5-HT_2A_ receptor - use of NAADP antagonist. (A) Schematic representation of signalling pathways associated with the 5-HT_2A_ receptor and pathway inhibitors. (B) Effect of 5-HT or 25B-NBOMe (10 µM) on NAADP concentrations in C6 cells compared to vehicle (**p=0.0011, **p=0.0053, one-way ANOVA with Dunnett’s test). (C) 5-HT concentration-dependent Ca^2+^ responses in the presence of Ned-19 (10 - 100 µM) in C6 cells. (D) As with (C) but for ATP. (E) As with (C) but with Ned-19 (10 - 100 µM) co-added. (F) As with (C) but in SH-SY5Y cells. (G) As with (C) but for 25B-NBOMe. (H) Summary graph showing efficacy of 5-HT_2A_ receptor agonists in C6 cells with and without Ned-19 (and Ca^2+^ store inhibitors - data taken from Fig. 1) (***p=0.0006, ****p<0.0001, ***p=0.0007, ***p=0.0002, Extra-sum-of-squares F-test with Dunnett’s test) (I) Summary graph showing efficacy of 5-HT with and without Ned-19 in SH-SY5Y cells (**p=0.0059, *p=0.037, *p=0.046, Extra-sum-of-squares F-test with Dunnett’s test). Each point or bar is the mean ± SEM value of five independent experiments performed in duplicates. Responses are relative to 10 µM 5-HT.

Next, we investigated the effect of the membrane-permeant NAADP antagonist Ned-19, which is thought to antagonise NAADP-regulation of TPCs^41^. Pre-treatment of C6 cells with Ned-19 (0.1 - 100 μM) caused a concentration-dependent decrease in the Ca**^2+^** response to 5-HT (0.1 nM - 10 μM) whilst having no effect on ATP (10 nM - 1 mM) (Fig. 2C, D; Suppl. Table 1,3). Ned-19 did not act via direct antagonism of the 5-HT_2A_ receptor because the drug (100 μM and 10 μM) had no effect on the 5-HT evoked Ca^2+^ response when added simultaneously with 5- HT (Fig. 2E; Suppl. Table 1). This contrasts with previous data reporting complete blockade of the 5-HT response by the selective 5-HT_2A_ receptor antagonist, MDL-100,907^28^. The inhibition of 5-HT by Ned-19 pretreatment was also seen in transfected SH-SY5Y cells (Fig. 2F; Suppl. Table 2). Moreover, in C6 cells, pretreatment with Ned-19 inhibited the Ca^2+^ response to the selective 5-HT_2A_ receptor agonist 25B-NBOMe, providing further evidence that the 5-HT_2A_ receptor activates a NAADP/lysosomal signalling pathway (Fig. 2G; Suppl. Table 4).

### NAADP signalling plays a role in Ca^2+^ response to psychedelic drugs

Given this link between the 5-HT_2A_ receptor and NAADP/lysosomal signalling, further experiments tested whether activation of this pathway was a feature of psychedelic drugs more generally. We have previously demonstrated that psilocin, 5-MeO-DMT and DOI evoked 5-HT_2A_ receptor mediated Ca^2+^ responses^28^. Pre-treatment of C6 cells with Ned-19 (100 μM) significantly reduced the magnitude of the Ca^2+^ response to each of these agonists (all at 0.1nM -10 μM) (Fig. 3A-C; Suppl. Table 4). It was of interest to know whether, compared to psychedelic drugs, newly emerging non-psychedelic 5-HT_2A_ receptor agonists might differentially affect NAADP/lysosomal signalling. However, as previously observed^28^, in C6 cells the non-psychedelic 5-HT_2A_ receptor agonist lisuride exhibited low efficacy in the form of a very small Ca^2+^ response, making it difficult to determine inhibition by Ned-19 (Fig. 3D; Suppl. Table 4; but see later).

**Figure 3.**
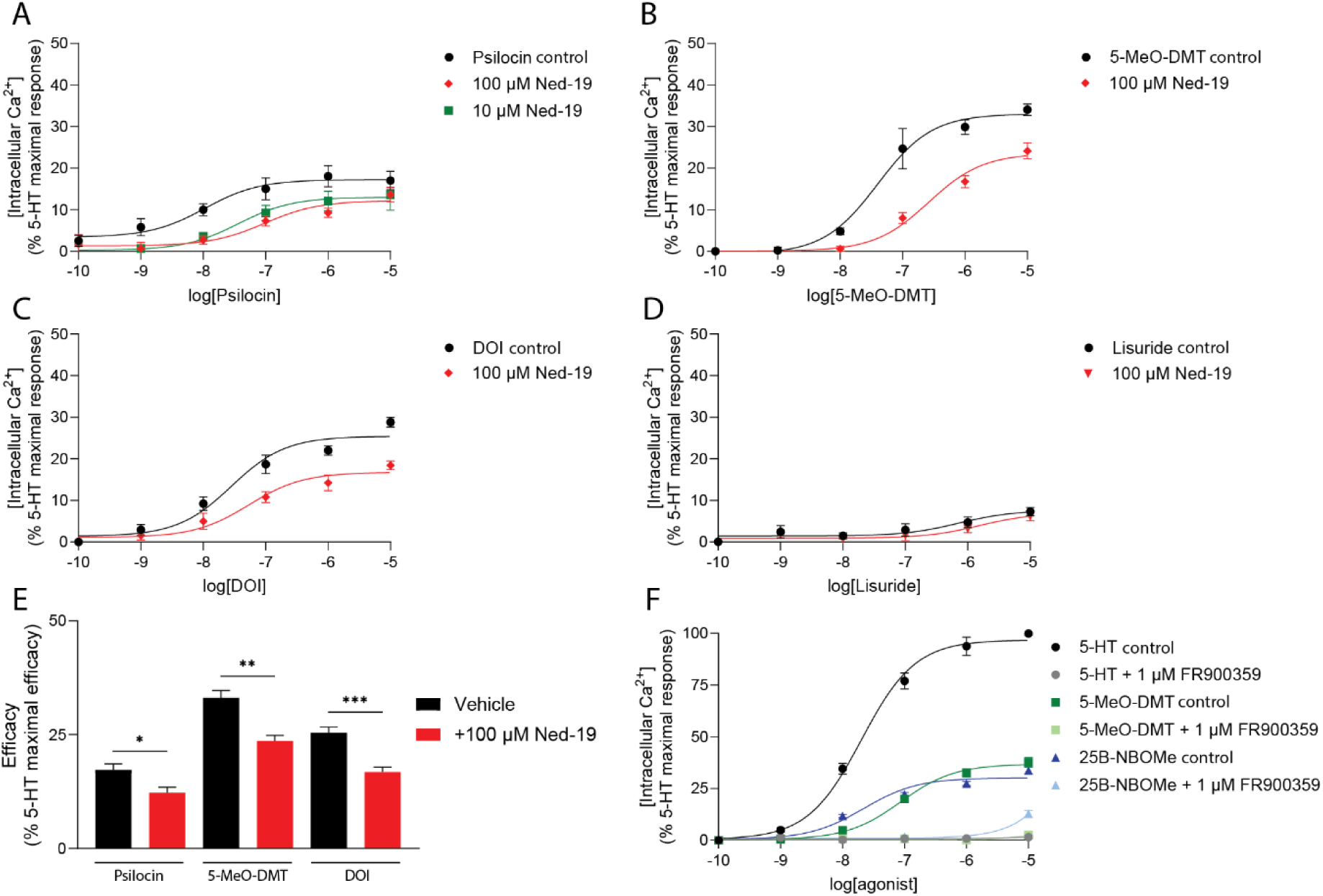
The NAADP/lysosomal pathway contributes to Ca^2+^ responses evoked by psychedelic drugs. Ca^2+^ responses in C6 cells to (A) psilocin, (B) 5-MeO-DMT, (C) DOI, (D) lisuride with and without Ned-19 (10 - 100 µM). (E) Summary graph showing efficacy of psychedelic drugs with and without Ned-19 in C6 cells (*p=0.0359, **p=0.0022, ***p=0.0008, Extra-sum-of-squares F-test with Dunnett’s test). (F) Ca^2+^ responses in C6 cells to 5-HT, 25B-NBOMe and 5-MeO-DMT with and without G_q_ inhibitor FR900359 (1 µM). Each point or bar is the mean ± SEM value of five independent experiments performed in duplicates. Responses are relative to 10 µM 5-HT.

### G_q_ coupling is required for 5-HT_2A_ receptor-mediated lysosomal Ca^2+^ release

We next investigated the mechanism by which the 5-HT_2A_ receptor might couple to NAADP generation and lysosomal Ca^2+^ release. Precisely how GPCRs mobilise NAADP formation and signalling is currently unclear^42^, although a previous study reported that cholecystokinin receptors require G_q_ coupling to evoke NAADP-dependent lysosomal Ca^2+^ release^43^. Here, the G_q_ inhibitor FR900359 was used to assess whether G_q_ activation was necessary for 5-HT_2A_ receptor-mediated lysosomal Ca^2+^ release. Pretreatment with FR900359 (1 µM) abolished the Ca^2+^ responses to each of 5-HT, 5-MeO-DMT and 25B-NBOMe (Fig. 3F), suggesting a requirement for G_q_ coupling in the lysosomal (plus ER) Ca^2+^ release in response to 5-HT_2A_ receptor activation.

### High resolution biosensor tracking confirms 5-HT_2A_ receptor-mediated mobilisation of lysosomal Ca^2+^ stores

To measure release of Ca^2+^ from lysosomes directly at single cell resolution, we employed a genetically-encoded Ca^2+^ indicator (G-GECO1.2) tethered to TPC2 (TPC2-G-GECO1.2^44^) (Fig. 4A). In individual C6 cells expressing TPC2-G-GECO1.2, 5-HT (10 µM) evoked Ca^2+^ responses and this effect was blocked by pre-treatment with MDL-100,907 (1 µM), demonstrating that the response was 5-HT_2A_ receptor-mediated (Fig. 4B, C). To control for the possibility that the lysosomal Ca^2+^ response signal measurement to 5-HT was contaminated by cytosolic Ca^2+^ spillover, a cytosolic version of the Ca^2+^ indicator (cytosolic G-GECO1.2) was also employed. Importantly, cells expressing lysosomal TPC2-G-GECO1.2 generated a significantly greater Ca^2+^ response to 5-HT than cells expressing cytosolic G-GECO1.2 (Fig. 4C), thereby directly demonstrating the 5-HT_2A_ receptor-induced mobilisation of Ca^2+^ from lysosomal stores. In confirmation of the latter, individual C6 cells expressing TPC2-G-GECO1.2 demonstrated a stronger Ca^2+^ response to 25B-NBOMe (10 µM) versus cytosolic G-GECO1.2, whereas ATP, which mobilised Ca^2+^ by a wholly IP_3_-linked mechanism, had no significant effect (Fig. 4D). Interestingly, lisuride (10 µM) elicited a response detectable by the TPC2-G-GECO1.2 biosensor, suggesting that even non-hallucinogenic 5-HT_2A_ receptor agonists are able to mobilise lysosomal Ca^2+^.

**Figure 4.**
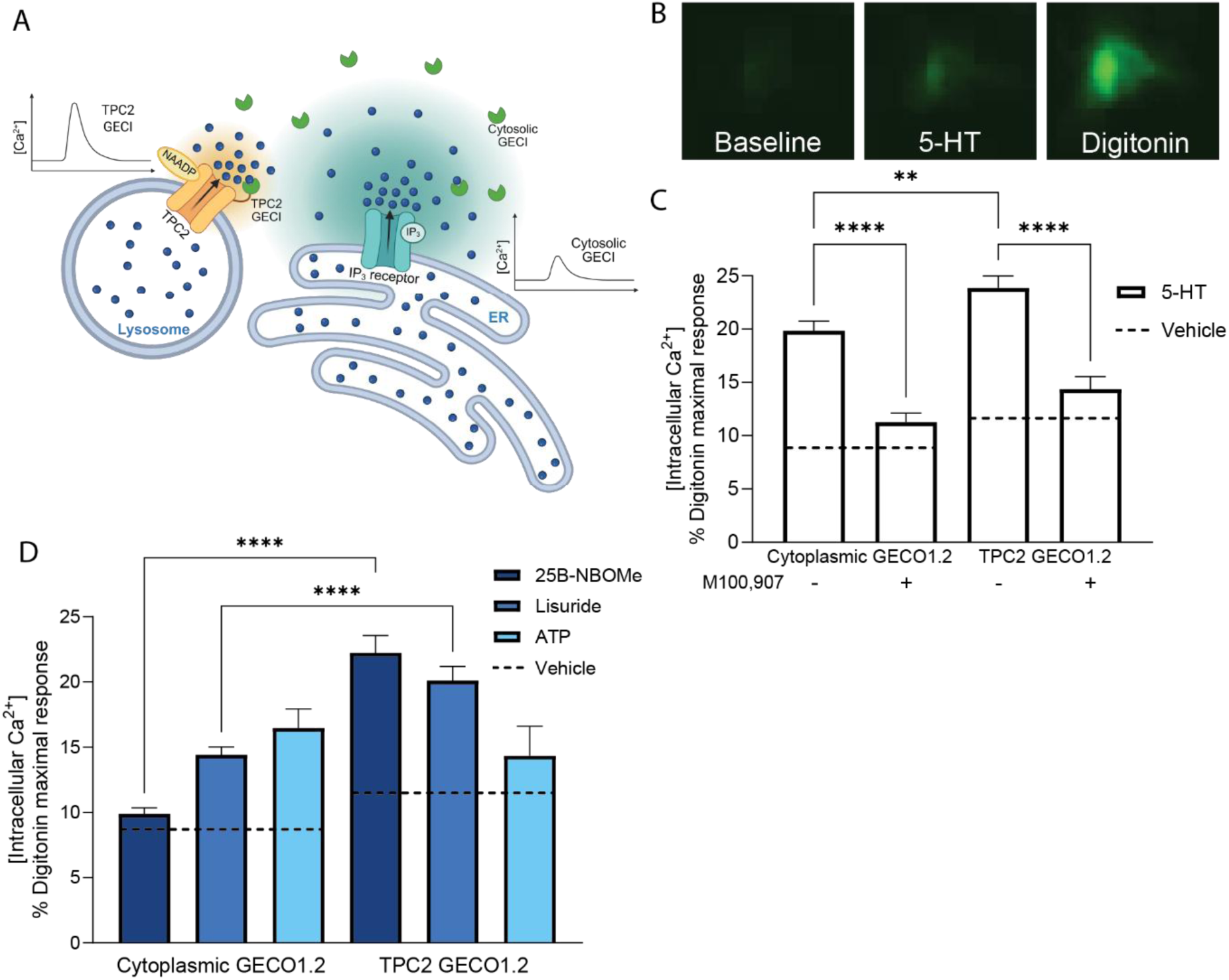
5-HT_2A_ receptor agonists release lysosomal Ca^2+^ - use of lysosomal Ca^2+^ tracker. (A) Schematic representation of TPC2 and cytosolic genetic calcium indicators (GECIs). (B) Representative images of a single cell transfected with TPC2-GECI at baseline (left), and after 5-HT (10 µM) (middle) or digitonin (75 µg/ml) (right). (C) Effect of 5-HT_2A_ receptor antagonist MDL-100,907 (1 µM) on Ca^2+^ responses to 5-HT (10 µM), measured by TPC2-or cytoplasmic-linked GECI in C6 cells (****p>0.0001, **p=0.0015, two-way ANOVA with Šidák’s test). (D) Ca^2+^ responses to 25B-NBOMe, Lisuride, ATP (10 µM), measured by TPC2-or cytoplasmic-linked GECI in C6 cells (****p<0.0001, one-way ANOVA with Šidák’s test). Bars represent mean ± SEM value of 3-5 independent experiments, n = 200-500 cells. Bars represent the mean ± SEM value of five independent experiments performed in duplicates.

### NAADP signalling plays a role in psychedelic drug-induced behavioural and neuroplasticity effects

Given the above evidence that psychedelic drugs activated NAADP/lysosomal signalling, the contribution of this pathway to behavioural and neuroplasticity effects of such agents in mouse models was investigated (Fig. 5A). Initial experiments examined the mouse head-twitch response which is widely accepted as a proxy for hallucinogenic effects of psychedelic drugs^7^. To this end, the head-twitch response to systemic administration of psilocin (1 mg/kg i.p.) was tested in mice with genetic knockout (KO) of two-pore channel TPC2, which is a NAADP target located on lysosomal membranes and required for NAADP-evoked Ca²⁺ release from lysosomal stores^45^. In TPC2 KO mice, the number of psilocin-induced head-twitches was reduced by ∼33% compared to wild-type controls (Fig. 5B), particularly during the first 8 min post-drug time interval (Fig 5C; Suppl. Table 5).

**Figure 5.**
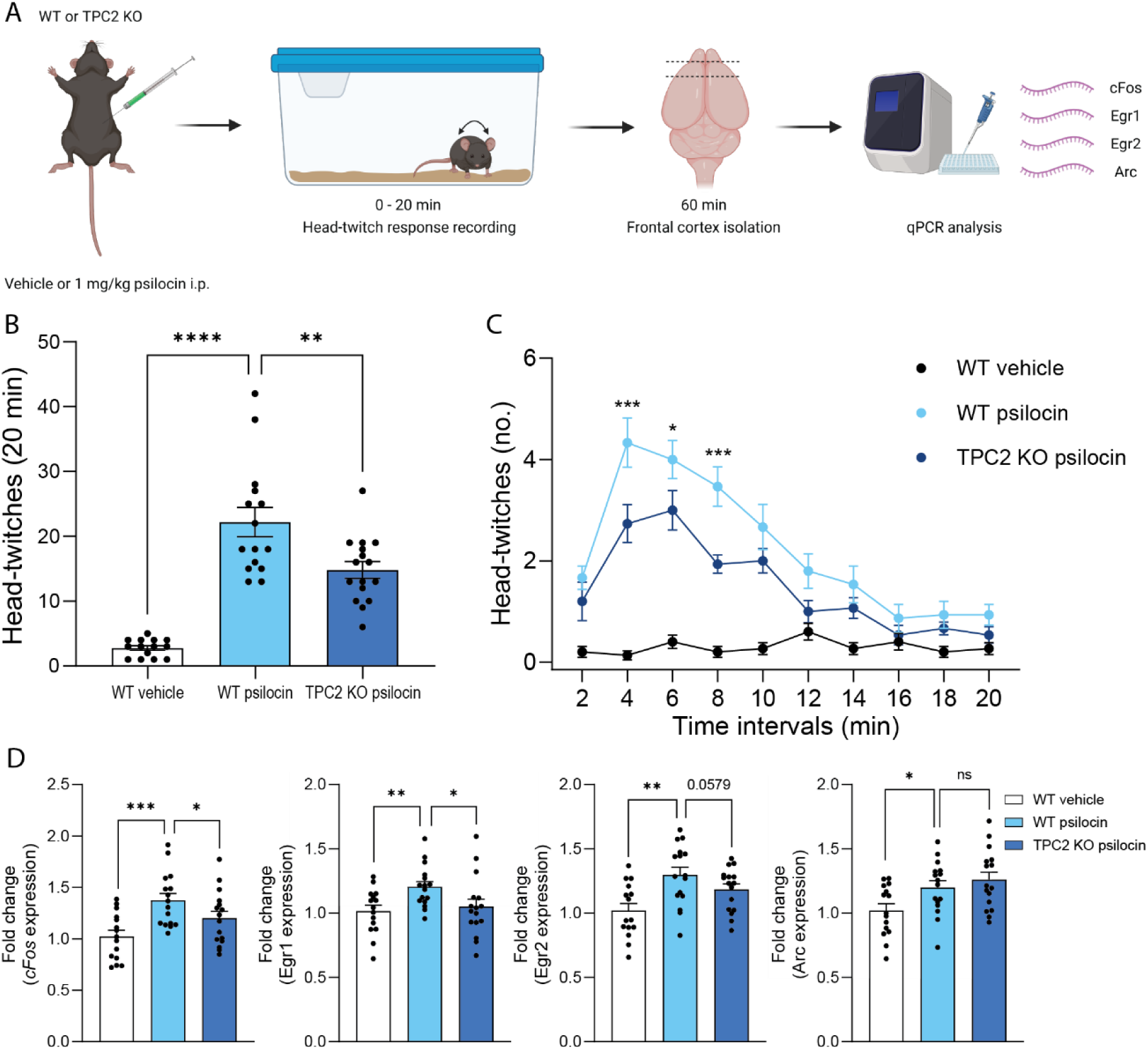
TPC2 knockout reduces psilocin-evoked head-twitches and cortical expression of plasticity-associated genes in mice. (A) In vivo experimentation workflow. (B) Effect of psilocin (1 mg/kg) or vehicle on total number of head-twitches in 20 min in TPC2 KO mice or WT controls (****p<0.0001, **p=0.00847, one-way ANOVA with Šidák’s test) (C) Effect of psilocin (1 mg/kg) or vehicle on the time-course of head-twitches in TPC2 KO mice or WT controls (***p=0.0003, *p=0.0248, ***p=0.0005, two-way ANOVA with post-hoc Tukey’s test) (D) Effect of psilocin (1 mg/kg) or vehicle on mRNA expression of *c-Fos* (***p=0.000353, *p=0.0188, one-way ANOVA with Šidák’s test)*, Egr-1* (**p=0.00449, *p=0.0353), *Egr-2* (**p=0.00184) and *Arc* (*p=0.0182) in frontal cortex of TPC2 KO mice or WT controls. Bars represent mean ± SEM, n = 15-16 mice.

Finally, we utilised TPC2 KO mice to test whether NAADP/lysosomal signalling also contributed to psychedelic drug-induced neuroplasticity using, as a marker, expression of a panel of plasticity-related genes in mouse cortex. In wildtype mice, systemic administration of psilocin (1 mg/kg i.p.) increased expression of each of *cFos*, *Egr-1*, *Egr-2* and *Arc* in frontal cortex compared to vehicle (Fig. 5D). In TPC2 KO mice this effect of psilocin was overall less than observed in wildtype mice (*c-fos*, *Egr-1* and *Egr-2* but not *Arc*; Fig 5D).

The validity of the TPC2 KO model was confirmed through qPCR analysis. Thus, cortical TPC2 mRNA was reduced to barely detectable levels in TPC2 KO versus wildtype mice (Suppl. Fig. 1A) whereas levels of G_q_ and 5-HT_2A_ receptor mRNA were unchanged (Suppl. Fig. 1B/C).

## Discussion

The 5-HT_2A_ receptor is strongly linked to the neuropharmacological and therapeutic effects of psychedelic drugs. It is generally assumed that GPCRs such as the 5-HT_2A_ receptor which utilise G_q_, generate important downstream signalling effects via increased formation of the second messenger IP_3_ and its mobilisation of Ca^2+^ stored in the ER. Here we investigated our finding that the 5-HT_2A_ receptor also has the capacity to mobilise Ca^2+^ from a non-ER store. Through this, we discovered the first evidence of a 5-HT_2A_ receptor-coupled signalling pathway involving the second messenger NAADP and mobilisation of lysosomal Ca^2+^ stores. Further, using mice lacking the lysosomal Ca^2+^ channel and NAADP target, TPC2, we found that TPC2 action was linked to the hallucinogenic and neuroplastic effects of psychedelic drugs.

Thus, in cells expressing the rat 5-HT_2A_ receptor (rat C6 glioma), 5-HT application increased formation of NAADP and evoked a rise in cytosolic Ca^2+^ that was reduced by depletion of lysosomal Ca^2+^ stores (bafilomycin) and the presence of a selective NAADP antagonist (Ned-19). Moreover, experiments using a selective biosensor to measure lysosomal Ca^2+^ (TPC2-G-GECO1.2) also detected a specific lysosomal Ca^2+^ response to 5-HT. Importantly, our finding of a 5-HT_2A_ receptor-coupled NAADP/lysosomal signalling pathway could be replicated in a second cell line expressing the human 5-HT_2A_ receptor (human SH-SY5Y neuroblastoma). Moreover, the pathway was activated by a range of psychedelic drugs with 5-HT_2A_ receptor agonist activity, including psilocin and 5-Me-O-DMT. The Ca^2+^ response to ATP lacked involvement of lysosomal stores, despite ATP likely acting via a purinergic G_q_-coupled receptor (P_2Y_), emphasising the specificity of the effect to 5-HT_2A_ receptor activation. It is presently unclear why 5-HT_2A_ and not P_2Y_ receptors couple to NAADP-mediated Ca^2+^ release but one possible explanation is that diversity in G_q_ protein βγ subunits^43^ confers diversity in signalling outcomes.

The size of the 5-HT_2A_ receptor-mediated lysosomal Ca^2+^ response was small relative to that of the ER Ca^2+^ response, however our data suggests that it is sufficiently important to trigger behavioural and neuroplasticity effects associated with 5-HT_2A_ receptor activation. Thus, in mice with a genetic loss of TPC2, a receptor which normally mediates NAADP-evoked release of Ca^2+^ from lysosomal stores, we observed a quantitative reduction in both behavioural and gene expression responses to the psychedelic agent, psilocin. Despite the lysosomal Ca^2+^ response being small, a previous study provided evidence that local TPC2-mediated Ca^2+^ release but not global Ca^2+^ release was responsible for macrophage phagocytosis^46^. It is also plausible that there is synergism between lysosomal and ER Ca^2+^ stores that results in an enhanced signalling response to 5-HT_2A_ receptor agonists and subsequent downstream functional effects. Indeed, previous reports suggest that a small local NAADP-stimulated Ca²⁺ signal can act as trigger for ER Ca^2+^ release, and that ER Ca^2+^ can similarly potentiate lysosomal Ca^2+^ release to eventually lead to larger global Ca²⁺ transients^47,48^. Thus, a combined ER and lysosome Ca^2+^ response may ultimately determine the specificity of the functional effects resulting from 5-HT_2A_ receptor activation.

It is currently uncertain which 5-HT_2A_ receptor signalling pathways are linked to psychedelic drug effects, although recent data favours a key role for G_q_ protein-versus β-arrestin-coupled pathways^22,28,29^. Here, 5-HT_2A_ receptor-mediated NAADP/lysosomal signalling was blocked by the G_q_ protein inhibitor FR900359, indicating that G_q_ protein-coupling is required for NAADP formation. Moreover, our data indicates that this NAADP/lysosomal signalling is necessary for behavioural and gene expression effects of psychedelic drug administration. Thus, TPC2 KO reduced 5-HT_2A_ receptor agonist-induced head-twitches, which are thought to be a surrogate marker of the hallucinogenic effects of these drugs. Evidence has suggested that this psychedelic experience is important for the antidepressant effect of these drugs^9^. Similarly, TPC2 KO reduced 5-HT_2A_ receptor agonist-induced expression of a panel of genes linked to neuroplasticity, a mechanism which is also considered important for the antidepressant properties of psychedelics as well as other therapeutic effects of these drugs^10,11^. In support of a role for NAADP/lysosomal signalling in neuroplasticity, mGluR1 receptor-stimulated long-term potentiation (LTP) and LTP-evoked dendritic spine growth were both reported to involve lysosomal Ca^2+^ release mediated by NAADP^49–52^.

In the context of the role of NAADP/lysosomal signalling in the therapeutic effects of psychedelic drugs, it is interesting that emerging evidence supports a plausible role for lysosomal biology in the pathophysiology of depression. Thus, there is molecular evidence of impaired lysosomal function in a rodent stress-based model of depression^53^ and reports of dysregulated autophagy, a key lysosomal-dependent mechanism, in postmortem brains of both animal models and depressed humans^53–55^. Moreover, evidence has suggested that multiple antidepressants, including the rapid acting antidepressant ketamine, exert antidepressant effects via induction of autophagy^56,57^. Given that TPC2-mediated lysosomal Ca^2+^ release has previously been reported to stimulate autophagy, it is plausible that 5-HT_2A_ receptor NAADP/lysosomal signalling could underlie antidepressant effects of psychedelic drugs^58,59^.

The current data provide insights into the therapeutic potential of future agents targeting NAADP/lysosomal signalling. The development of 5-HT_2A_ receptor agonists with a bias towards NAADP/lysosomal signalling merits consideration. In this regard it is of interest that we found that the non-hallucinogenic 5-HT_2A_ receptor agonist tested (lisuride) also elicited NAADP/lysosomal signalling but had lower Ca^2+^ signalling efficacy overall than hallucinogenic agonists (psilocin, 5-MeO-DMT). It is plausible that drugs which directly target NAADP/lysosomal signalling might be a starting point for the future drug development. It is noteworthy that there is a hint of symmetry between the functional effects of 5-HT_2A_ and TPC2 receptor ligands. Thus, the TPC2 antagonist tetrandrine is reported to inhibit 5-HT_2_ receptor agonist-evoked head-twitches^60^, as observed here with TPC2 KO. Although there currently exist few TPC2 receptor agonists, one such agonist has recently shown therapeutic potential in cellular models of Parkinson’s Disease^61^, which accords with reports that 5-HT_2A_ receptor agonists may have neuroprotective properties^62^. Thus, greater understanding of TPC2 neuropharmacology and agonist development in the context of therapeutics certainly warrants further exploration.

## Methods

### Cell culture

Rat C6 glioma cells expressing the endogenous 5-HT_2A_ receptor^63,64^ (ATCC CCL-107, RRID:CVCL_0194) were maintained in Ham’s F12 nutrient mix containing 2 mM Glutamax, 10% (v/v) FBS and 100 IU μg^−1^ ml^−1^ penicillin/streptomycin. Human SH-SY5Y neuroblastoma cells (RRID:CVCL_0019) transfected with the human 5-HT_2A_ receptor^65^ were maintained in culture medium: Dulbecco’s Modified Eagle Medium (DMEM) containing 2 mM Glutamax, 10% (v/v) foetal bovine serum (FBS), 100 IU μg^−1^ ml^−1^ penicillin/ streptomycin and 480-μg ml^−1^ G418 (to maintain transfection selection pressure). Cells were grown at 37°C in a humidified atmosphere of 95% air and 5% CO_2_. Plate reader Ca^2+^ assays were carried out on both C6 or SH-SY5Y cells while single-cell Ca^2+^ measurements and NAADP enzyme-linked immunosorbent assay (ELISA) measurements were obtained in C6 cells only.

### Plate reader assay of agonist-evoked cytosolic Ca^2+^

Measurements of agonist-evoked cytosolic Ca^2+^ levels in C6 and SH-SY5Y cells were carried out as previously described^28^. Baseline fluorescence was measured on a plate reader (BMG Optima) every 3 s for 30 s prior to addition of one of 10 μl agonist, after which fluorescence was recorded for a further 3 min (a time frame which allowed for the agonist-induced response to reach a maximum). In each well, the final concentration of DMSO was 0.1% (v/v).

In some experiments, before application of 5-HT or 5-HT_2A_ receptor agonist (typically 0.1 nM - 10 µM), cells were pre-treated with; i) the Ca^2+^-ATPase inhibitor thapsigargin (1 µM for 60 min) to assess ER Ca^2+^ contribution to the response, ii) the H^+^-ATPase inhibitor bafilomycin (1 µM for 45 min) to assess lysosomal Ca^2+^ contribution to the response, or iii) the NAADP antagonist Ned-19 (0.1 µM - 100 µM for 15 min) to assess the role of NAADP. Concentration-response curves were generated as described previously^28^.

### Single-cell measurements of agonist-evoked cytosolic and lysosomal Ca^2+^

C6 cells were kept in antibiotic-free cell culture medium for 60 min prior to transfection with DNA encoding either TPC2-G-GECO1.2 or cytosolic G-GECO1.2 (200 ng/well) using Viafect transfection reagent diluted in Opti-MEM (Gibco) at a 1:5 ratio. Then, 10 µl of transfection mix was added to each well of a 96-well black/clear bottom plate before addition of ∼20,000 cells diluted in 100 µl of culture medium (antibiotic-free, 10% dialysed FBS).

After 48 h C6 cells were washed with HBSS before addition of 90 μl HBSS. Cells were then imaged on a fluorescent microscope (Invitrogen Evos FL Auto 2) at 20X magnification with images captured every 0.07s. Cells were imaged for 30s prior to addition of 10 µl 5-HT or 5-HT agonist (10 µM) and further recording for 2 min, then finally addition of 25 µl digitonin (75 µg/ml) and 25 µl CaCl_2_ (2 M) and recording for a further 2 min. The 2 min post-drug period allowed for both the agonist- and digitonin-induced responses to reach a maximum. In some experiments before stimulation with 5-HT (10 µM), cells were pre-treated for 15 min with the selective 5-HT_2A_ receptor agonist, MDL-100,907 (1 µM).

Images were analysed in ImageJ^66^ using an adaptation of a previously used protocol^67^. Briefly, regions of interest were drawn to delineate individual cells before plotting z-axis profiles of fluorescence intensity across time for each cell. Fluorescence increase for each individual cell was calculated as follows:

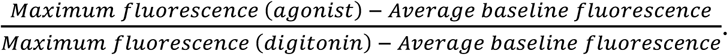

### ELISA measurement NAADP levels in C6 cell extracts

Levels of NAADP were measured in C6 cell extracts using a competitive ELISA assay (St John’s Laboratories). Cells were cultured in T175 flasks to supply 6 x 10^6^ cells per well for assay using ELISA. On the day of experiment cells were removed from the flask using trypsin (Gibco), centrifuged and then resuspended and aliquoted into Eppendorf tubes (one Eppendorf for each well of the assay). An equal volume of 5-HT, 25B-NBOMe or vehicle was then added and mixed thoroughly before immediately adding an equal volume of ice-cold 1.5 M HClO_4_ to stop the reaction. NAADP was extracted as previously described^68^. Briefly, cells were sonicated, incubated on ice for 15 min, then centrifuged at 9000g for 10 min to pellet the denatured protein. The acid supernatant was then neutralised with an equal volume of 2 M KHCO_3_, then the sample was centrifuged at 9000g for 10 min to remove the KClO_4_ precipitate, leaving only the NAADP extract.

The ELISA assay was carried out according to manufacturer’s instructions: 50 µl of NAADP extract sample and 50 µl biotin-labelled NAADP analogue was added to each well and incubated for 1 h at 37°C. Wells were then washed three times with wash buffer, followed by addition of 100 µl of avidin-conjugated HRP and incubation for 1 h at 37°C. Wells were again washed three times, then 90 µl of substrate reagent was added before incubation at 37°C for 20 min. Finally, 50 µl of stop solution was added to each well. Absorbance at 450nm was then immediately measured using the SPECTROstar Omega plate reader and concentration of NAADP was determined using a calibration curve generated according to manufacturer’s instructions.

### Drugs

Psilocin (supplied by Compass Pathways), 5-MeO-DMT (Cambridge Bioscience), DOI HCl (Cambridge Bioscience), 25B-NBOMe HCl (Chiron), Ned-19 (Bio-techne), thapsigargin (Abcam), bafilomycin (Alfa Aesar) and MDL-100,907 (Tocris) were all dissolved in DMSO (Sigma Aldrich) whereas 5-HT HCl (Enzo Life Sciences) was dissolved in deionised water, to obtain 10 mM stock solutions. Ned-19 (Tocris) was dissolved in DMSO to obtain a 100 mM stock solution. On the day of experiment, working solutions of drugs were obtained by serial dilutions with HBSS.

### Animals

Mice were group housed (4-6 per cage) in individually ventilated cages in a temperature-controlled room (21 °C) on a 12-h light/dark cycle. Mice had ad libitum access to food and water, and cages were lined with sawdust bedding and contained cage enrichment (sizzle nest and cardboard tubes). Experiments were conducted in the light phase. Three days prior to experiments, all mice were habituated to handling, a scruff-hold for injection, and exposure to the experimental room.

Experiments were conducted on C57BL/6J wildtype mice and TPC2 KO (Tpcn2Gt(YHD437)Byg) mice^45^ (C57Bl/6J background, gift from Prof Francis Platt, Dept of Pharmacology, Oxford), with groups being a mix of males and females (aged 8/9 weeks). Experiments followed the UK animals (Scientific Procedures) act of 1986 and ARRIVE guidelines.

### Measurement of mouse head-twitches

On the day of experiment, mice were habituated to the experimental room and their individual cages for 90 min before injection. Immediately after i.p. administration of psilocin or vehicle, mice were placed individually in clear cages and video recorded for 20 min. Head-twitch counting was performed post-experiment by two observers blind to treatment and genotype, and the counts average were taken.

### qPCR analysis

After cervical dislocation, the whole brain was removed, snap frozen in isopentane and stored at -80°C. The frontal cortex was dissected, and RNA extraction was performed using the TRIzol method (Rio et al., 2010) (Qiagen RNeasy Mini Kit, Qiagen). cDNA conversion and subsequent qPCR were performed as described previously (Radford-Smith et al., 2022). For each sample, 800 ng of RNA was converted to cDNA using a high-capacity cDNA Reverse Transcription Kit (Life Technologies) and T100 Thermocycler (Bio-Rad). The primer sequences used are as follows: cFos forward: AGGGGCAAAGTAGAGCAGCTA, cFos reverse: CAATCTCAGTCTGCAACGCA (PrimerBank ID 31560587b2); Egr1 forward: TTGTGGCCTGAACCCCTTTT, Egr1 reverse: AGATGGGACTGCTGTCGTTG^69^; Egr2 forward: TGCTAGCCCTTTCCGTTGA, Egr2 reverse: TCTTTTCCGCTGTCCTCGAT^70^; Arc forward: GGAGGGAGGTCTTCTACCGTC, Arc reverse: CCCCCACACCTACAGAGACA (PrimerBank ID 86604725c2); Tpcn2 forward: TAAAGTACCGCTCCATCTACCA, Tpcn2 reverse: GCAGACGTTCGAGTAATACCAG^71^; Gnaq forward: TTGGTTCGAGAGGTTGATGTG, Gnaq reverse: CGTCGTCTGTCGTAGCACTC (Primerbank ID 145966786c2); Htr2a forward: CAGGCAAGTCACAGGATAGC, Htr2a reverse: TTAAGCAGAAAGAAAATCCCACAG^72^.

For qPCR, a 384-well plate was used with 10 µl reaction volume per well, 20 ng cDNA, 6 pmol forward and reverse primers, 1.3 μl nuclease-free water and 5 μl Precision®PLUS qPCR Master Mix with SYBRgreen (Primer Design). The qPCR protocol consisted of enzyme activation at 95°C for 2 min, 40 cycles of 95°C for 10 s and 60°C for 1 min, then holding at 4°C. Each sample was run in duplicate and values averaged.

qPCR data was analysed using the ΔCt method^73,74^. Outliers were identified using ROUT analysis (GraphPad Prism) and these samples were excluded from further analysis.

### Statistical analysis

Data sets were tested for normality using the Shapiro-Wilk test (GraphPad Prism, version 10.4.1). For the concentration-response curves, differences between efficacy (E_max_) and potency (pEC_50_) values in the presence and absence of inhibitors was statistically tested using Extra sum-of-squares F-tests. This test compares the goodness-of-fit of two alternative nested models; one in which E_max_ and pEC_50_ are shared between curves in the presence or absence of an inhibitor, and one in which E_max_ and pEC_50_ are different between these curves. The null hypothesis is that the simpler model (E_max_ and pEC_50_ are shared between datasets) is correct and thus treatment with an inhibitor has no effect on the concentration-response curve of the agonist alone. If the F-test was significant, post-hoc Dunnett’s tests (Suppl. Tables 1-3) or two-tailed t-tests (Suppl. Table 4) were used to assess differences between individual E_max_ and pEC_50_ values. For the single-cell Ca^2+^ measurements, data were analysed with a two-way (for 5-HT with or without MDL-100,907) or one-way ANOVA and where this was significant, followed by post-hoc Sidak’s Multiple Comparisons tests. For the NAADP ELISA assay, data were analysed with a one-way ANOVA and where this was significant, followed by post-hoc Dunnett’s Multiple Comparisons tests.

RStudio (RStudio 2024.04.1) was used to perform statistical analysis on the data. The lmer package, part of the lme4 package, was used to perform linear mixed-effects modelling, with sex and experimental cohort treated as covariates, as previously described^75^. The lmer function was followed by Type III ANOVA with Satterthwaite’s method for estimating the degrees of freedom. If significance was reached using ANOVA, the analysis was followed by post-hoc Tukey’s Honest Significant Difference testing. Data are presented as mean ± SEM values with individual values plotted. P<0.05 was considered statistically significant.

## Author Contributions

*Conceptualisation*: Aurelija Ippolito, Trevor Sharp, Grant Churchill. *Data curation*, *Formal analysis, Investigation*: Aurelija Ippolito. *Methodology*: Aurelija Ippolito, Antony Galione, Grant Churchill, Trevor Sharp. *Writing—original draft*: Aurelija Ippolito, Trevor Sharp. *Review and editing*: Antony Galione, Trevor Sharp, Gary Gilmour, Grant Churchill. *Supervision*: Trevor Sharp, Grant Churchill, Gary Gilmour. *Resources*: Gary Gilmour. *Funding acquisition*: Trevor Sharp

## Acknowledgements

This study was supported by a Medical Research Council iCASE studentship (Aurelija Ippolito) in association with Compass Pathways Plc.

## Data Availability

The data that support the findings of this study are available from the corresponding author upon reasonable request.

