## Supplementary tables and figures for "A novel 5-HT_2A_ receptor-linked signalling pathway involving NAADP-mediated lysosomal Ca^2+^ release"

**Supplementary Table 1 | Summary of $Ca^{2+}$ responses to 5-HT in the presence of Ned-19 (pre-treatment or co-addition), bafilomycin or thapsigargin in C6 cells.** $E_{max}$ and $pEC_{50}$ values are the mean ± SEM values of five independent experiments performed in duplicates. $E_{max}$ values are relative to 10 µM 5-HT. Extra sum-of-squares F-tests compared the efficacy and potency of the concentration-response curves of 5-HT alone with those in the presence of 0.1 – 100 µM Ned-19, 1 µM bafilomycin or 1 µM thapsigargin. Post-hoc Dunnett's tests (p-values shown here) then compared $E_{max}$ and $pEC_{50}$ values in the presence of inhibitors to those of 5-HT alone. *p<0.05, **p<0.01, ***p<0.001, ****p<0.0001, ns = not significant in F-test so no post-hoc testing was performed.

| Agonist | Pre-treatment | $E_{max}$ ± SEM | p-value | $pEC_{50}$ ± SEM | p-value |
|---|---|---|---|---|---|
| 5-HT | None | 100.81 ± 2.70 | N/A | 7.54 ± 0.073 | N/A |
| 5-HT | 100 µM Ned-19 | 75.05 ± 3.71 | 0.0007*** | 6.96 ± 0.13 | 0.0288* |
| 5-HT | 10 µM Ned-19 | 71.58 ± 4.26 | 0.0002*** | 7.50 ± 0.20 | 0.9991 |
| 5-HT | 1 µM Ned-19 | 97.27 ± 4.58 | 0.9214 | 7.28 ± 0.14 | 0.4935 |
| 5-HT | 0.1 µM Ned-19 | 99.46 ± 4.42 | 0.9975 | 7.33 ± 0.13 | 0.6655 |
| 5-HT | 1 µM Bafilomycin | 82.15 ± 2.54 | 0.0006*** | 7.28 ± 0.090 | 0.77 |
| 5-HT | 1 µM Thapsigargin | 14.20 ± 2.77 | <0.0001**** | 6.48 ± 0.500 | 0.0469* |
| 5-HT | None | 106.03 ± 5.13 | N/A | 7.43 ± 0.15 | N/A |
| 5-HT | 100 µM Ned-19 (co-addition) | 105.73 ± 2.85 | ns | 7.22 ± 0.075 | ns |
| 5-HT | 10 µM Ned-19 (co-addition) | 108.26 ± 3.88 | ns | 7.47 ± 0.11 | ns |

**Supplementary Table 2 | Summary of $Ca^{2+}$ responses to 5-HT in the presence and absence of Ned-19, bafilomycin or thapsigargin pre-treatment in SH-SY5Y cells.** $E_{max}$ and $pEC_{50}$ values are the mean ± SEM values of five independent experiments performed in duplicates. $E_{max}$ values are relative to 10 µM 5-HT. Extra sum-of-squares F-tests compared the efficacy and potency of the concentration-response curves of 5-HT alone with those in the presence of 0.1 – 100 µM Ned-19, 1 µM bafilomycin or 1 µM thapsigargin. Post-hoc Dunnett's tests (p-values shown here) then compared $E_{max}$ and $EC_{50}$ values in the presence of inhibitors to those of 5-HT alone. *p<0.05, **p<0.01, ***p<0.001, ****p<0.0001.

| Agonist | Pre-treatment | $E_{max}$ ± SEM | p-value | $pEC_{50}$ ± SEM | p-value |
|---|---|---|---|---|---|
| 5-HT | None | 98.87 ± 2.51 | N/A | 7.99 ± 0.084 | N/A |
| 5-HT | 100 µM Ned-19 | 78.02 ± 5.36 | 0.0059** | 6.81 ± 0.18 | <0.0001**** |
| 5-HT | 10 µM Ned-19 | 82.56 ± 3.79 | 0.0337* | 7.93 ± 0.15 | 0.9964 |
| 5-HT | 1 µM Ned-19 | 83.41 ± 4.57 | 0.046* | 7.82 ± 0.18 | 0.8411 |
| 5-HT | 0.1 µM Ned-19 | 94.22 ± 3.50 | 0.833 | 8.08 ± 0.13 | 0.9776 |
| 5-HT | 1 µM Bafilomycin | 84.66 ± 3.11 | 0.0201* | 7.99 ± 0.12 | >0.9999 |
| 5-HT | 1 µM Thapsigargin | 24.12 ± 4.19 | <0.0001**** | 7.99 ± 0.60 | >0.9999 |

**Supplementary Table 3 | Summary of $Ca^{2+}$ responses to ATP in the presence and absence of Ned-19, bafilomycin or thapsigargin pre-treatment in C6 cells.** $E_{max}$ and $pEC_{50}$ values are the mean ± SEM values of five independent experiments performed in duplicates. $E_{max}$ values are relative to 10 µM ATP. Extra sum-of-squares F-tests compared the efficacy and potency of the concentration-response curves of ATP alone with those in the presence of 0.1 – 100 µM Ned-19, 1 µM bafilomycin or 1 µM thapsigargin. Post-hoc Dunnett's tests (p-values shown here) then compared $E_{max}$ and $pEC_{50}$ values in the presence of inhibitors to those of ATP alone. *p<0.05, **p<0.01, ***p<0.001, ****p<0.0001. N.D. = no detectable response.

| Agonist | Pre-treatment | $E_{max}$ ± SEM | p-value | $pEC_{50}$ ± SEM | p-value |
|---|---|---|---|---|---|
| ATP | None | 116.55 ± 4.34 | N/A | 5.26 ± 0.10 | N/A |
| ATP | 100 µM Ned-19 | 114.70 ± 4.54 | 0.8468 | 5.09 ± 0.15 | 0.7936 |
| ATP | 10 µM Ned-19 | 110.27 ± 5.07 | 0.8468 | 5.33 ± 0.13 | 0.9946 |
| ATP | 1 µM Ned-19 | 107.32 ± 4.51 | 0.5483 | 5.25 ± 0.11 | >0.9999 |
| ATP | 0.1 µM Ned-19 | 108.35 ± 5.08 | 0.6579 | 5.34 ± 0.13 | 0.9907 |
| ATP | 1 µM Bafilomycin | 123.22 ± 5.96 | 0.8123 | 5.02 ± 0.12 | 0.5284 |
| ATP | 1 µM Thapsigargin | N.D. | <0.0001**** | N.D. | N/A |

**Supplementary Table 4 | Summary of $Ca^{2+}$ responses to 5-$HT_{2A}$ receptor agonists in the presence and absence of Ned-19, bafilomycin or thapsigargin pre-treatment in C6 cells.** $E_{max}$ and $pEC_{50}$ values are the mean ± SEM values of five independent experiments performed in duplicates. $E_{max}$ values are relative to 10 µM 5-HT. Extra sum-of-squares F-tests compared the efficacy and potency of the concentration-response curves of 5-$HT_{2A}$ receptor agonists alone with those in the presence of 0.1 – 100 µM Ned-19, 1 µM bafilomycin or 1 µM thapsigargin. Post-hoc Dunnett's tests or two-tailed t-tests (p-values shown here) then compared $E_{max}$ and $pEC_{50}$ values in the presence of inhibitors to those of the 5-$HT_{2A}$ receptor agonist alone. *p<0.05, **p<0.01, ***p<0.001, ****p<0.0001, ns = not significant in F-test so no post-hoc testing was performed. †= $E_{max}$ value derived from incomplete concentration-response curve. ‡ = $pEC_{50}$ value is an estimate due to low $E_{max}$.

| Agonist | Pre-treatment | $E_{max}$ ± SEM | p-value | $pEC_{50}$ ± SEM | p-value |
|---|---|---|---|---|---|
| Psilocin | None | 17.25 ± 1.31 | N/A | 7.97 ± 0.31 | N/A |
| Psilocin | 100 µM Ned-19 | 12.21 ± 1.21 | 0.0359* | 7.01 ± 0.28 | 0.0854 |
| Psilocin | 10 µM Ned-19 | 13.04 ± 1.44 | 0.079 | 7.44 ± 0.34 | 0.3913 |
| Psilocin | 1 µM Bafilomycin | 8.10 ± 0.70 | <0.0001**** | 7.77 ± 0.27 | 0.9905 |
| Psilocin | 1 µM Thapsigargin | N.D. | <0.0001**** | N.D. | 0.66 |
| 5-MeO-DMT | None | 33.06 ± 1.64 | N/A | 7.40 ± 0.14 | N/A |
| 5-MeO-DMT | 100 µM Ned-19 | 23.60 ± 1.27 | 0.0022** | 6.58 ± 0.13 | 0.0028** |
| DOI | None | 25.45 ± 1.20 | N/A | 7.55 ± 0.15 | N/A |
| DOI | 100 µM Ned-19 | 16.77 ± 1.12 | 0.0008*** | 7.26 ± 0.20 | 0.2982 |
| 25B-NBOMe | None | 48.55 ± 4.53 | N/A | 8.23 ± 0.36 | N/A |
| 25B-NBOMe | 100 µM Ned-19 | 22.25 ± 0.78 | 0.0039** | 8.14 ± 0.17 | 0.825 |
| Lisuride | None | 7.61 ± 1.46 | N/A | 6.14 ± 0.45 | N/A |
| Lisuride | 100 µM Ned-19 | 7.03 ± 1.39 | ns | 5.81 ± 0.37 | ns |
| TBG | None | 42.63 ± 4.07† | N/A | 4.56 ± 0.14‡ | N/A |
| TBG | 100 µM Ned-19 | 56.26 ± 22.61† | 0.5829 | 3.94 ± 0.32‡ | 0.1402 |

**Supplementary Table 5 | Post-hoc analysis of the effect of genotype on time-course data of the head-twitch response for wildtype and TPC2 KO mice dosed with psilocin.** Given the significant genotype-timepoint interaction, post-hoc Tukey's tests (p-values shown here) were carried out comparing WT vs KO mice dosed with 1 mg/kg psilocin within each 2-minute time interval. *p<0.05, **p<0.01, ***p<0.001. Group sizes are as in Fig. 5C.

| Time interval (min) | p-value |
|---|---|
| 0 – 2 | 0.366 |
| 2 – 4 | 0.0003*** |
| 4 – 6 | 0.0248* |
| 6 – 8 | 0.0005*** |
| 8 – 10 | 0.142 |
| 10 – 12 | 0.0752 |
| 12 – 14 | 0.317 |
| 14 – 16 | 0.492 |
| 16 – 18 | 0.595 |
| 18 – 20 | 0.399 |

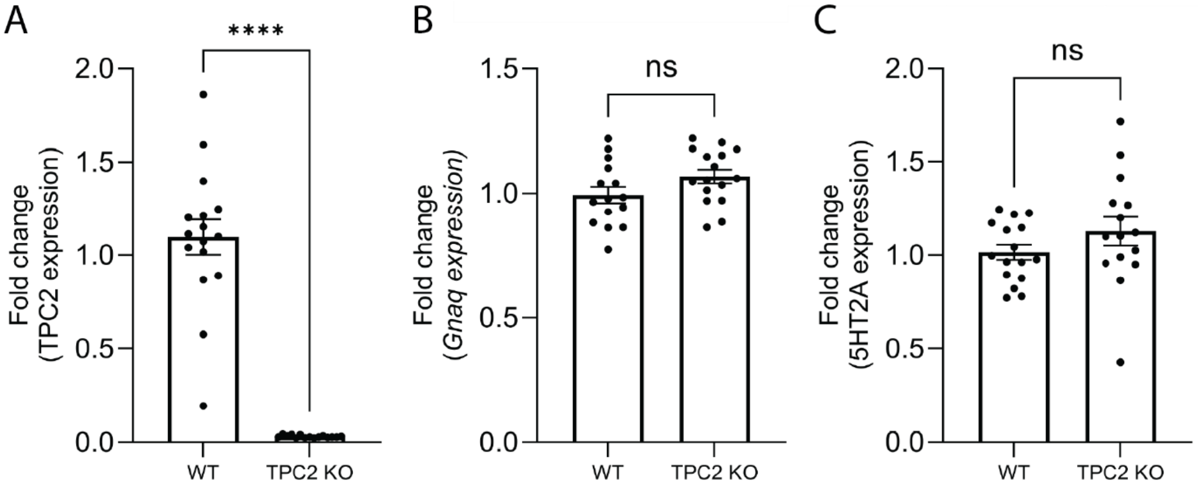


**Supplementary Figure 1 | Effect of TPC2 KO on mRNA expression of TPC2, $G_q$ and the 5-$HT_{2A}$ receptor.** (A) TPC2 (B) $G_q$ and (C) 5-$HT_{2A}$ receptor mRNA in the frontal cortex of WT and TPC2 KO mice (n=15-16). Columns represent mean ± SEM values, with individual values indicated by symbols. Data were analysed by an unpaired t-tests.****p<0.0001, ns = not significant.
